# Loss of tomato geranylgeranyl diphosphate synthase 2 increases monoterpenoid levels and enhances immune responses to bacterial infection

**DOI:** 10.1101/2025.03.24.644926

**Authors:** Julia Pérez-Pérez, Miguel Ezquerro, Sooyeon Lim, Sun-Hwa Ha, Mª Pilar López-Gresa, Manuel Rodríguez-Concepción, Purificación Lisón

## Abstract

Many plastidial isoprenoids, including diterpenes and photosynthesis-related isoprenoids such as carotenoids and chlorophylls, derive from C20 geranylgeranyl diphosphate (GGPP), produced by GGPP synthase (GGPPS) enzymes. Heterodimers of GGPPS and non-catalytic type I small subunit (SSU-I) proteins produce C10 geranyl diphosphate (GPP), the precursor of monoterpenes.

Three plastidial GGPPS isoforms, referred to as SlG1-3, are present in tomato (*Solanum lycopersicum*). Here we explored their contribution to the production of volatile organic compounds (VOCs) of isoprenoid origin under normal conditions and in response to infection with *Pseudomonas syringae* pathovar *tomato* (*Pst*).

Edited lines lacking SlG2 showed a distinctive VOC profile compared to unedited (WT) plants and mutants impaired in SlG1 or SlG3. In particular, only *slg2* mutants showed constitutively increased levels of GPP-derived hydroxylated monoterpenes (HMTPs). Upon *Pst* infection, *slg2* plants accumulated higher levels of salicylic acid (SA) and exhibited increase resistance compared to WT controls, resulting in reduced levels of VOCs associated to cell death.

Our findings suggest that SlG2 regulates GPP synthesis, potentially by specifically competing with other GGPPS isoforms for heterodimerization with SSU-I. Increased GPP production in *slg2* plants could lead to higher HMTPs levels, which may result in elevated SA content, and subsequently enhanced protection against bacterial infection.

## INTRODUCTION

Isoprenoids,also known as terpenoids, represent one of the most diverse groups of plant metabolites, consisting of a wide variety of molecules synthesized from the 5-carbon (C5) units isopentenyl diphosphate (IPP) and dimethylallyl diphosphate (DMAPP). IPP and DMAPP are synthesized by the mevalonic acid (MVA) pathway in the cytosol or the methylerythritol 4-phosphate (MEP) pathway in plastids (Vranová et al., 2013; Rodríguez-Concepción & Boronat, 2015). Condensation of one or more molecules of IPP to one molecule of DMAPP by short-chain prenyltransferases generates prenyl diphosphates of increasing number of carbons, such as C10 geranyl diphosphate (GPP), C15 farnesyl diphosphate (FPP), and C20 geranylgeranyl diphosphate (GGPP). These prenyl diphosphates are the precursors to produce the main groups of isoprenoids.

Most plastidial isoprenoids derive either from GPP or GGPP. The main GPP-derived products are monoterpenes, mostly involved in defense against biotic stress (Zhou & Pichersky, 2020; Perez-Perez et al., 2024). A much higher diversity of isoprenoids derives from plastidial GGPP, including defensive diterpenes and photosynthesis-related terpenoids such as carotenoids, prenylated quinones (phylloquinone, plastoquinone), and chlorophylls (Barja & Rodriguez-Concepcion, 2021). Plastid-localized GGPP synthase (GGPPS) enzymes producing GGPP from MEP-derived IPP and DMAPP have been found to form homo and heterodimers. Typically, GGPPS homodimers and heterodimers of different GGPPS paralogs synthesize GGPP whereas heterodimers with catalytically inactive members of the GGPPS-like small subunit (SSU) family show altered enzymatic properties. While heterodimerization with type I SSU proteins (SSU-I) mostly produces GPP, heterodimerization with type II SSU (SSU-II) proteins enhance GGPP production (Zhou & Pichersky, 2020; Barja & Rodriguez-Concepcion, 2021; Song et al., 2023). GPP can also be synthesized by homodimeric GPP synthases (GPPS), although it is generally assumed that most GPP for monoterpene synthesis originates from heterodimeric GGPPS/SSU-I enzymes (Tholl et al., 2004; Chen et al., 2015; Hivert et al., 2020; Zhou & Pichersky, 2020; Song et al., 2023).

Three genes encode plastidial GGPPS enzymes in tomato (*Solanum lycopersicum*), here referred to as SlG1, SlG2 and SlG3 (Barja et al., 2021). SlG1 (Solyc11g011240) is involved in the production of diterpenes in leaves and strigolactones in roots (Ament et al., 2006; Ezquerro et al., 2023). SlG3 (Solyc02g085700) is the main housekeeping enzyme providing GGPP to produce photosynthesis-related isoprenoids (chlorophylls and carotenoids) in tomato, and it also contributes to diterpene production in response to bacterial infection (Barja et al., 2021; Ezquerro et al., 2023; Burbano-Erazo et al., 2024). SlG2 (Solyc04g079960) appears to assist SlG3 when extra production of carotenoids is required in different tissues but it does not appear to participate in leaf diterpene synthesis (Barja et al., 2021; Ezquerro et al., 2023; Burbano-Erazo et al., 2024). Consistent with these relative contributions in providing GGPP, tomato CRISPR mutants defective in the SIG3 isoform exhibit stronger phenotypes related to reduced carotenoid levels (e.g., paler leaves and fruits) compared to those deficient in SlG2, whereas SlG1-defective plants are almost undistinguishable from wild-type controls (Barja et al., 2021; Ezquerro et al., 2023; Burbano-Erazo et al., 2024). The contribution of plastidial GGPPS isoforms in the synthesis of plastidial isoprenoids, aside from diterpenes and photosynthetic pigments, remains still unexplored. In particular, little is known about their role in the production of defense-related volatile organic compounds (VOCs) beyond diterpenes.

VOCs have important roles for defense and plant-to-plant communication (Yu et al., 2024). A non-targeted GC-MS analysis of VOCs emitted by Rio Grande tomato plants, carrying the *Pseudomonas syringae* pathovar *tomato* (*Pst*) resistance gene (*Pto*), after infection with an avirulent strain of the bacterium revealed that the specific aroma emitted by resistant plants included hydroxylated monoterpenes (HMTPs) such as α-terpineol, 4- terpineol, and linalool (López-Gresa et al., 2017). HMTPs were later shown to confer resistance to bacteria by immune signaling involving both the salicylic acid (SA) and the MEP pathway intermediate methylerythritol cyclodiphosphate (MEcPP) (Pérez-Pérez et al., 2024). The VOCs associated with successful infection -referred to as the scent of death- in tomato cultivars such as MicroTom, which lack the *Pto* resistance gene and are therefore unable to resist bacterial infection, remain unidentified.

Here we aimed to investigate the contribution of tomato GGPPS paralogs to the production of VOCs emitted by MicroTom plants under normal conditions and after infection with *Pst*. By comparing the VOC profile of previously generated CRISPR lines defective in SlG1, SlG2 or SlG3 with the unedited MicroTom parental we found that only *slg2* mutants showed a differential profile in the absence of bacterial infection. Strikingly, HMTPs were increased in the volatilome of SlG2-defective plants, leading to increased levels of SA, but not MEcPP, and enhanced resistance to bacterial infection. Furthermore, we identified VOCs associated with the scent of death in the *Pst*-MicroTom infection and confirmed that their levels were reduced in *Pst*-*slg2* tomato plants.

## MATERIAL AND METHODS

### Plant material and growth conditions

Tomato (*Solanum lycopersicum* L.) var. MicroTom plants and homozygous CRISPR-edited mutant alleles defective in GGPPS isoforms SlG1 (*slg1-1*), SlG2 (*slg2-1*) or SlG3 (*slg3-1*) (Barja et al., 2021; Ezquerro et al., 2023) were used for the experiments. For sterilization, we used a mixture of sodium hypochlorite and distilled water (1:1) with sequential washes of 5, 10, and 15 min to completely remove the hypochlorite. Germinated seeds were placed in 12-cm-diameter pots with vermiculite and peat and transferred to a climate-controlled growth chamber. Growth conditions were approximately 50% relative humidity, 26°C of temperature, and a photoperiod of 14h of light and 10h of darkness. VOCs and other metabolites were measured in leaflet samples from the second, third and fourth true leaves of 4-week-old tomato plants. Samples were snap-frozen in liquid nitrogen immediately after collection.

### Infection assays

The bacterial strain utilized in this research was *Peudomonas syringae* pv. *tomato* DC3000 (*Pst*) (Ntoukakis et al., 2009). The methods for bacterial cultivation and plant inoculation followed those outlined by López-Gresa et al. (2018). In brief, 4- week-old tomato plants were inoculated through immersion in a solution containing 0.05% Silwet L-77 and a bacterial suspension IN 10 mM MgCl_2_ with an optical density at 600 nm of 0.1. For mock inoculations, plants were dipped in 10 mM of MgCl_2_ solution with 0.05% Silwet L-77 without the bacterial inoculum. To measure colony-forming units (cfu), three leaf disks (1 cm² each) from the second, third and fourth true leaves were collected and ground, and serial dilutions of the infected tissue were plated on King’s B agar medium containing 50 mg/L rifampicin. The cfu were counted after 48h of incubation at 28°C.

### VOC analysis

Frozen leaf powder (100 mg) was placed in a 10-mL headspace vial with a screw cap. A mixture of 1 mL saturated CaCl_2_ solution and 100 μL of 750 mM EDTA (pH 7.5, adjusted with NaOH) was added. The sample was gently mixed and then sonicated for 5 min. VOC extraction was performed using headspace solid-phase micro-extraction (HS-SPME) (Rambla et al., 2015). The sample underwent pre-incubation and extraction at 50°C for 10 and 20 min, respectively. A 65 μm DVB/PDMS fiber (Supelco, Bellefonte PA, United States) was used for adsorption, and desorption occurred in the gas chromatograph injection port for 1 min at 250°C in splitless mode. This process was automated with a CombiPAL autosampler (CTC Analytics, Zwingen, Switzerland). The separation of compounds was carried out using an Agilent 8860 gas chromatograph equipped with a DB-5 ms fused silica capillary column (60 m long, 0.25 mm i.d., 1 μm film thickness) as previously described by López-Gresa et al., (2017). The oven temperature program started at 40°C for 2 min, then increased by 5°C per min until reaching 250°C, and was held at 250°C for 5 min. Helium was used as the carrier gas at a steady flow rate of 1.2 mL/min. Detection was conducted with an Agilent 5977B mass spectrometer in EI mode (70 eV ionization energy; source temperature 230°C). Data was acquired in scan mode (m/z range 35–250; six scans per second). The Enhanced MassHunter software (Agilent, Santa Clara CA, United States) was used for recording and processing chromatograms and mass spectra. To conduct an untargeted analysis of the VOC profile, the GC-MS data were processed using MetAlign software (Wageningen, Netherlands), which facilitated chromatogram alignment and quantification of each MS feature. The processed data set was then analyzed with PCA using SIMCA-P software (v. 11.0, Umetrics, Umeå, Sweden), applying unit variance (UV) scaling for the analysis. Several VOCs were conclusively identified by comparing their mass spectra and retention times with those of pure standards. All standards were sourced from Sigma–Aldrich (Madrid, Spain). Additionally, some compounds were tentatively identified by comparing their mass spectra with the NIST 05 Mass Spectral library, and these are marked with an asterisk. To quantify MeSA, a standard curve was constructed using known concentrations of the volatile compound to determine its concentration in ppm.

### SA and GA measurements

To extract salicylic acid (SA) and gentisic acid (GA), 0.25 g of frozen, homogenized leaf tissue were resuspended in methanol containing 25 mM *o*- anisic acid as an internal standard. The mixture was centrifuged for 10 min and sonicated for 10 min. The resulting supernatant was dried under a nitrogen stream. Identification and quantification was carried out with the method described by Vázquez Prol et al. (2021) using a calibration curve relative to the internal standard.

### JA measurements

To quantify JA, 250 mg of frozen leaf tissue were mixed with 80% methanol containing 1% acetic acid and dihydrojasmonate as the internal standard (OlChemim, Olomouc, Czechia). The mixture was shaken for 1 h at 4°C, then stored overnight at -20°C. After centrifugation, the supernatant was evaporated to dryness using a vacuum evaporator. The resulting residue was reconstituted in 1% acetic acid and purified through an Oasis HLB reverse-phase column (Seo et al., 2011). The dried eluate was re-dissolved in 5% acetonitrile with 1% acetic acid, and JA was analyzed using a reverse-phase Ultra Performance Liquid Chromatography (UPLC) system coupled to a Q-Exactive mass spectrometer with an Orbitrap detector (Thermo Fisher Scientific, Waltham MA, USA). Targeted Selected Ion Monitoring (SIM) was employed for detection. Chromatographic separation was performed on a 2.6 µm Accucore RP-MS column (50 mm × 2.1 mm i.d.; Thermo Fisher Scientific) using a gradient of 5–50% acetonitrile with 0.05% acetic acid as the solvent system at a flow rate of 400 µL/min over 14 min. JA concentrations were calculated using calibration curves prepared with an authentic JA standard (OlChemIm) and processed with Xcalibur 2.2 SP1 build 48 and TraceFinder software.

### MEcPP measurements

The quantification of MEcPP was conducted following the methodology outlined by Baidoo et al. (2014), with slight modifications. Frozen homogenized tissue (100 mg) was extracted using a 13 mM ammonium acetate buffer (pH 5.5). The resulting extract was dried under a stream of nitrogen gas and reconstituted in the UPLC mobile phase, consisting of 73% acetonitrile and 27% 50 mM ammonium carbonate in water (v/v). Analysis was carried out on an Orbitrap Exploris 120 mass spectrometer paired with a Vanquish UHPLC System (Thermo Fisher Scientific). Liquid chromatography employed reverse-phase ultraperformance liquid chromatography using a BEH Amide column (1.7 µm particle size, 2.1 × 150 mm; Waters Corp.). The samples were analyzed in isocratic mode for 14 min, with a flow rate of 0.2 mL/min and an injection volume of 5 µL. The column temperature was maintained at 30°C. Ionization was achieved via heated electrospray ionization (H-ESI) in positive mode, and data acquisition was performed in full scan mode with a resolution of 120,000 (measured at full width at half maximum). Methionine sulfone and D4-succinic acid served as internal standards. Calibration curves were generated using an MEcPP chemical standard (Echelon Biosciences, Salk Lake City UT, United States) to enable absolute quantification. Data processing was carried out using TraceFinder software (Thermo Scientific).

### RNA extraction and RT-qPCR

RNA extraction and cDNA synthesis from tomato leaves were performed using a silica membrane-based column kit (Macherey-Nagel GmbH, Dueren, Germany) following the manufacturer’s instructions. cDNA was synthesized from 1 µg of RNA using the PrimeScript RT reagent kit Perfect Real Time (Takara Bio Inc., Otsu, Shiga, Japan) according to the provided protocol. Quantitative PCR (qPCR) was conducted as previously described (Campos et al., 2014). Each reaction was carried out on a 96-well plate with a final volume of 10 µL. SYBR Green PCR Master Mix (Applied Biosystems) was used as the fluorescent marker, and the *actin ACT4* (Solyc04g011500) gene served as the endogenous reference. The qPCR primers for *SLG1* and *SLG2* were obtained from Barja et al., 2021, and those for *ICS* and *actin* from Pérez-Pérez et al., 2024. The remaining primers were designed using the Primer3 online tool (http://primer3.ut.ee/). The primer sequences are listed in the Suplementary Table S2.

### Computational analyses of protein structure and docking

Individual and heterodimeric protein structures of SlG1, SlG2, SlG3 and SSU-I were predicted using AlphaFold3 (Abramson et al., 2024) after retrieving protein sequences from the UniProt database and removing transit peptides to prevent interference with dimer formation. ChimeraX (Pettersen et al., 2021) was employed for structural visualization. PDBePISA (Krissinel & Henrick, 2007) was used to analyze interface characteristics including interface atom and residue counts, solvent-accessible area (SASA, Å²), solvation energy (kcal/mol), and the specific residues involved in complex formation. Docking simulations with AutoDock vina (Huey et al., 2012) predicted the binding positions of IPP and DMAPP substrates on the heterodimers. Moleonline server (Pravda et al., 2018) identified the active sites by selecting the channel surrounded by the substrate-binding region and the catalytic site predicted.

## RESULTS

### SlG2 Is a Key Regulator of Monoterpenoid Production in Tomato

To determine the possible contribution of individual GGPPS paralogs to the production of VOCs in tomato, we used previously generated CRISPR lines defective in SlG1 (allele *slg1-1*), SlG2 (allele *slg2-1*) or SlG3 (allele *slg3-1*) together with the unedited wild-type (WT) MicroTom parental (Barja et al., 2021; Ezquerro et al., 2023). Leaf samples from 4- week-old WT and mutant plants were employed for a non-targeted metabolomic analysis by GC-MS followed by a multivariate statistical study to assess their differential VOC content (see Materials and Methods).

The principal component analysis (PCA) illustrates the distinct clustering of the VOC profiles among WT and mutant tomato lines (*slg1*, *slg2*, and *slg3*). The first principal component (PC1), accounting for 35.2% of the variance, clearly separate the groups (Fig. 1). The *slg2* mutant forms a distinct cluster, markedly separated from the WT and other mutant lines along PC1, suggesting significant differences in VOC profiles. By contrast, *slg1* and *slg3* mutants overlap more closely with the WT, indicating more similar VOC profiles.

**Fig. 1.**
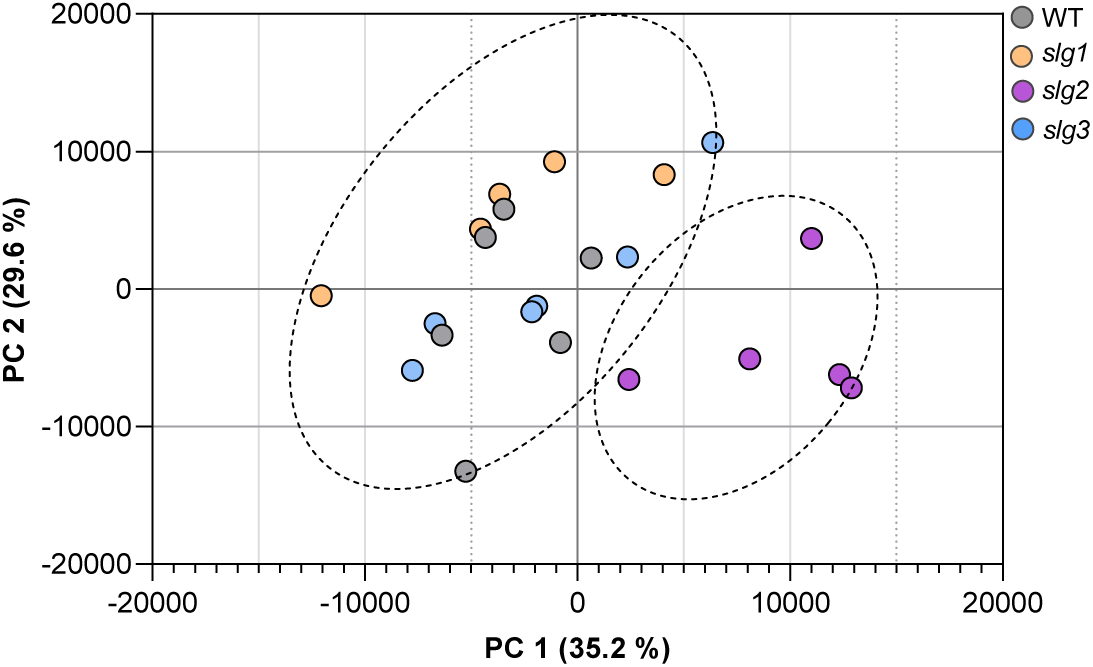
Principal component analysis (PCA) of VOC profiles unveils a differential volatilome for *sgl2* plants. Dots represent replicates for the indicated WT and mutant lines. Score plot of the PCA was based on the whole array of the mass spectra within a m/z range from 35 to 250. The dotted ellipses represent the 95% confidence intervals for each group.

To further investigate the impact of loss of SlG2 function in VOC production, we analyzed the differential compounds emitted by *slg2* mutants. A loading plot analysis revealed a notable increase in C10 monoterpenes, including β-myrcene, α-terpinene, trans-β- ocimene, D-limonene, and terpinolene, along with HMTPs such as β-terpineol, linalool, terpinene-4-ol, and α-terpineol in *slg2* mutants (Table 1). They were all unequivocally confirmed by using pure standards. The targeted quantification of some of these HMTPs confirmed that their levels were significantly higher in *slg2* (Fig. 2).

**Fig. 2.**
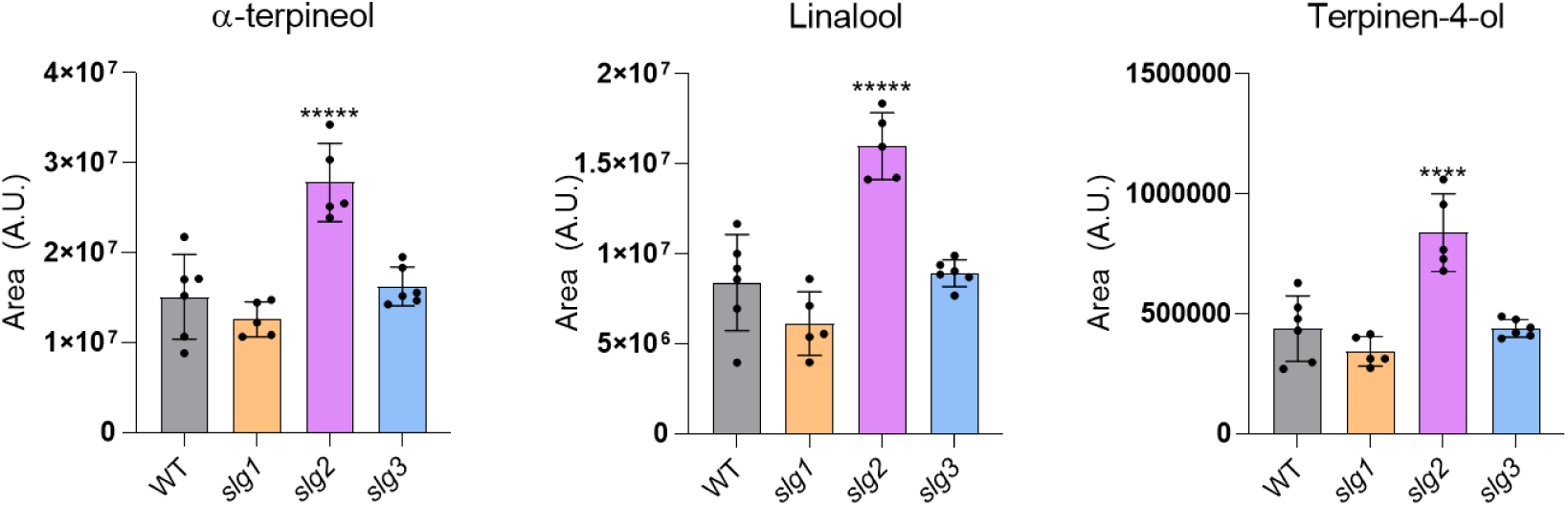
Mutant *sgl2* leaves release higher levels of HMTPs compared to WT, *slg1* and *slg3* lines. other mutants. The y axis represents arbitrary units (AU) corresponding to the quantification of the area under the chromatogram peak. Values represent the mean ± SD of five independent biological replicates (n = 5). Asterisks indicate statistically significant differences with the parental WT (one-way ANOVA with comparison test: ****, p <0.001; *****, p <0.0001).

**Table 1.** List of differentially emitted VOCs in *slg2* compared to WT samples. Only those showing a fold change (FC) >1.5 and p-values < 0.05 are shown. Family: MTP, monoterpenes; HMTP, hydroxylated monotepenes. RT, retention time. Ion, major ion fragments used for quantification.

| slg2 vs WT mock | VOC family | Ion | R.T (min) | Fold Change |
| --- | --- | --- | --- | --- |
| *β-Myrcene* | MTP | 93 | 23.12 | 1.89 |
| *α-Terpinene* | MTP | 121 | 24.49 | 1.91 |
| *trans-β-Ocimene* | MTP | 93 | 24.71 | 1.95 |
| *D-Limonene* | MTP | 68 | 24.95 | 1.78 |
| *β-Terpineol* | HMTP | 63 | 25.53 | 2.00 |
| *Terpinolene* | MTP | 93 | 26.90 | 1.91 |
| *Linalool* | HMTP | 93 | 27.01 | 1.77 |
| *Terpine-4-ol* | HMTP | 71 | 30.25 | 1.95 |
| *α-Terpineol* | HMTP | 59 | 30.61 | 1.65 |

### *In silico* analysis predicts a preferential interaction of SSU-I with SlG2 but a higher activity of the SlG3/SSU-I dimer

The most parsimonious explanation for increased levels of monoterpenes and HMTPs in the *slg2* mutant is an enhanced supply of their GPP precursor. To investigate whether the specific loss of SlG2 impacted the expression of genes encoding enzymes potentially altering GPP production, we carried out a RT-qPCR quantification of transcripts encoding the remaining GGPPS paralogs, SlG1 and SlG3, which can produce GPP by heterodimerization with SSU-I (Fig. 3). We also analyzed the level of transcripts encoding GPP synthase GPPS (Solyc08g023470) and the SSU proteins SSU-I (Solyc07g064660) and SSU-II (Solyc09g008920) in the same samples of WT and *slg2* leaves (Fig. 3).

**Fig. 3.**
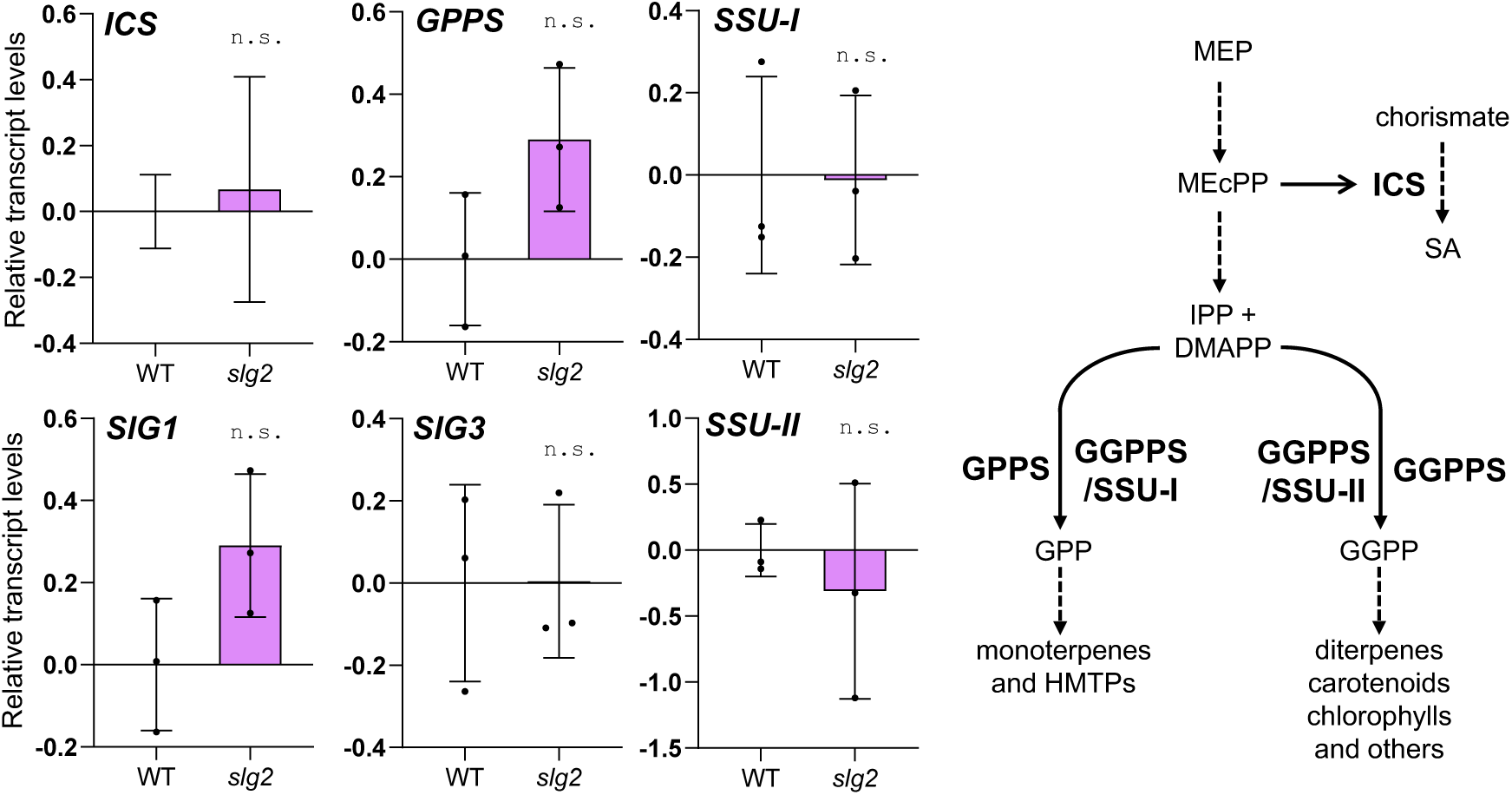
Expression of genes potentially impacting monoterpenoid or SA biosynthesis is similar in WT and *sgl2* leaves. Plots show RT-qPCR data on the levels of transcripts from the indicated genes, encoding enzymes that catalyze the steps indicated in the pathway cartoon. Transcript levels are shown relative to those in WT leaves, and they correspond to the mean and SD of n = 3 biological replicates. No statistically significant differences (n.s.) between WT and *slg2* samples were observed.

Although a trend towards higher levels of *GPPS* and *SlG1* transcripts was observed, no statistically relevant changes were found for any of the genes analyzed (Fig. 3).

Next, we used computational methods to investigate whether differential formation or/and activity of GPP-producing GGPPS/SSU-I heterodimers could explain the observed accumulation of monoterpenoids in SlG2-lacking plants (Fig. 4). The mint (*Mentha piperita*) GPPS adopts a heterotetrameric structure composed of two heterodimers, each formed by SSU and GGPPS-like large subunit (LSU) monomers (Chang et al., 2010; Hsieh et al. 2010). Substrate entry and product release is controlled by the AC-loop of the LSU subunit, which regulates the opening and closing of the active site where GPP is synthesized. In the SSU subunit, the R-loop regulates substrate entry and product release from LSU by interacting with the AC-loop (Chang et al., 2010; Hsieh et al. 2010). In tomato, the presence of the AC-loop in SlG1, SlG2, and SlG3, and the R-loop in SSU-I was predicted using sequence alignment and AlphaFold3 (Supplementary Fig. S1). The R-loop of the tomato SSU-I protein was surprisingly long compared to other plants homologs (Supplementary Fig. S1a), and it could potentially form β-sheet structures that might regulate the AC-loop of SlG1, SlG2, or SlG3 (Supplementary Fig. S1b). AlphaFold3 predicted that, in the SlG2/SSU-I heterodimer, the R-loop β-sheet structure adopted a conformation that obstructed the active site (Fig. 4 and Supplementary Fig. S1c). By contrast, the R-loop was predicted to be located away from the active site and to form no β-sheet structures in SlG1/SSU-I and SlG3/SSU-I heterodimers (Fig. 4 and Supplementary Fig. S1c), potentially causing a more flexible structure of the AC-loop and allowing, better ligand docking and product (GPP) release compared to the SlG2/SSU-I heterodimer.

**Fig. 4.**
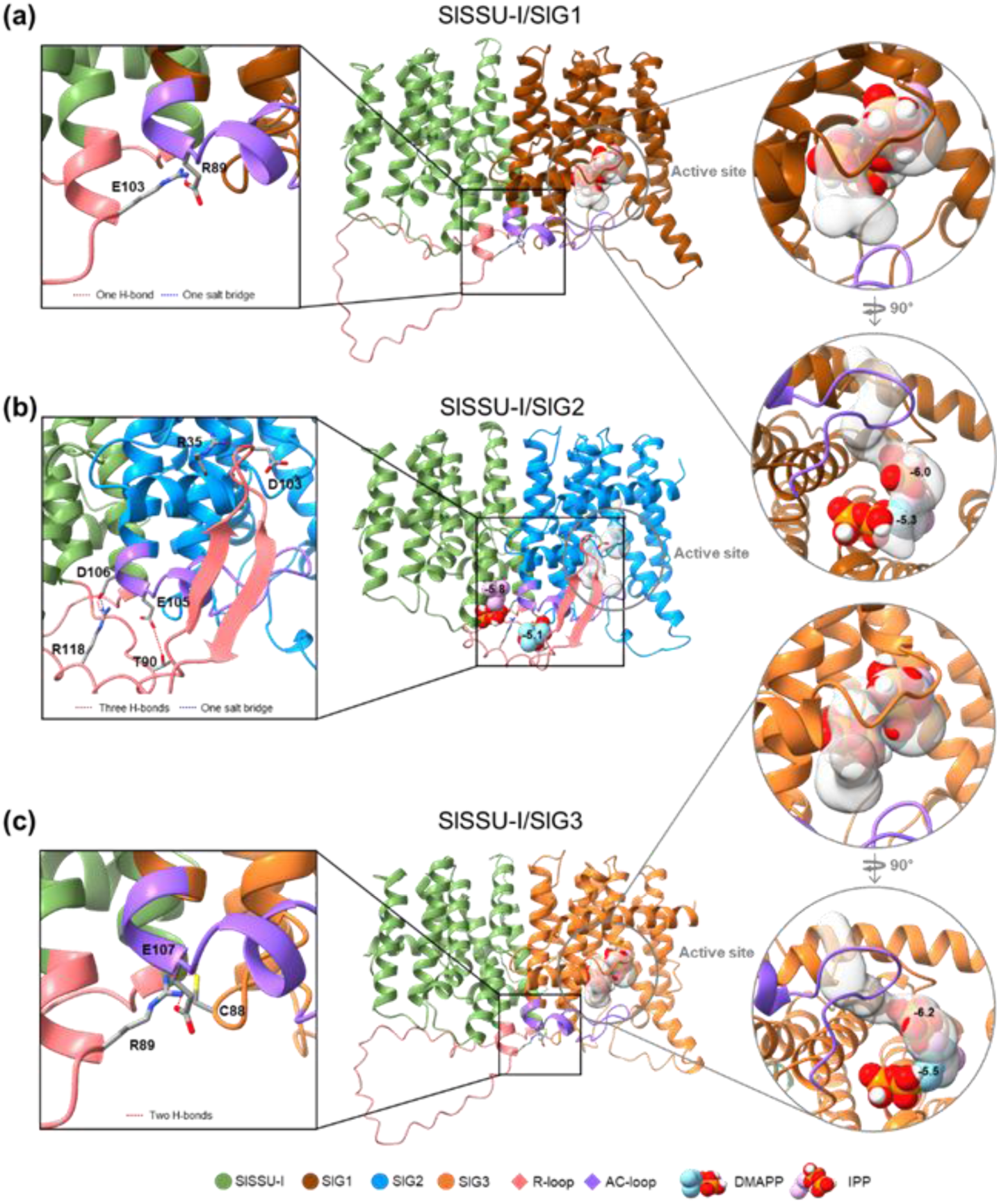
Structural prediction and docking analysis of tomato GGPPS/SSU-I heterodimers. Protein structures were predicted using AlphaFold3 and visualized with ChimeraX. H-bonds and salt bridges formed by residues within the R-loop were predicted using PDBePISA. Docking simulations of two substrates, IPP and DMAPP, were performed using AutoDock Vina (binding affinity in kcal/mol is shown as negative numbers). Active site and channel formation were predicted using MoleOnline server.

To further explore the potential differences in enzymatic activities, two additional algorithms for substrate docking and channel formation (AutoDock Vina and MoleOnline, respectively) were consecutively applied to the three heterodimeric protein-binding structures (Fig. 4). IPP and DMAPP substrates were properly located within the active site along the predicted channel in SlG1/SSU-I and SlG3/SSU-I, but not in SlG2/SSU-I (Fig. 4). Docking affinity for IPP and DMAPP was also higher in SlG1/SSU-I (-6.0 and -5.3) and SlG3/SSU-I (-6.2 and -5.5) than in SlG2/SSU-I (-5.8 and -5.1). These results strongly suggest that SlG2/SSU-I might be much less efficient than SlG1/SSU-I or SlG3/SSU-I in converting IPP and DMAPP into GPP.

We further analyzed the interface properties of the heterodimers and the specific residues involved in complex formation using PDBePISA. Strikingly, the SlG2/SSU-I interface was found to contain higher number of atoms and residues, a larger solvent-accessible surface area (Å²), and more negative free energy of solvation gained (kcal/mol) compared to SlG1/SSU-I and SlG3/SSU-I (Supplementary Table S1), suggesting the possibility of a stronger interaction of SSU-I with SlG2 than with SlG1 or SlG3. PDBePISA analysis of specific residues for chemical bonding, such as hydrogen bonds and salt bridges, further supported a greater interaction strength in the case of SlG2/SSU-I (Fig. 4).

Together, these comprehensive computational simulations predict that while tomato SSU-I likely binds more strongly to SlG2 than to SlG1 and SlG3, the SlG1/SSU-I and SlG3/SSU-I heterodimers might be more enzymatically efficient to produce GPP than SlG2/SSU-I. We conclude that monoterpenoid synthesis might rely on SlG2/SSU-I activity in WT plants, whereas loss of SlG2 would allow SlG1 or/and SlG3 to bind SSU-I and produce more GPP (and eventually HMTPs) in the *slg2* mutant.

### The tomato immune response to *Pst* is activated in SlG2-defective mutants

As previously mentioned, the involvement of HMTPs in defense against the bacterium *Pseudomonas syringae* DC3000 (*Pst*) has been well described in tomato (Lopez-Gresa et al., 2017; Pérez-Pérez et al., 2024). Moreover, there is also evidence of their involvement in resistance to *Pst* in *Arabidopsis thaliana* (Riedlmeier et al., 2017). We hypothesized that the increased levels of HMTPs observed in *slg2* plants might improve their resistance to *Pst*. To test this hypothesis, *slg2* and WT plants were infected with the bacterium *Pst* and bacterial counts were subsequently performed (Fig. 5). Indeed, *slg2* mutants exhibited significantly higher resistance to *Pst* infection compared to their parental WT line, as indicated by the markedly lower number of bacterial colonies (Fig. 5a).

**Fig. 5.**
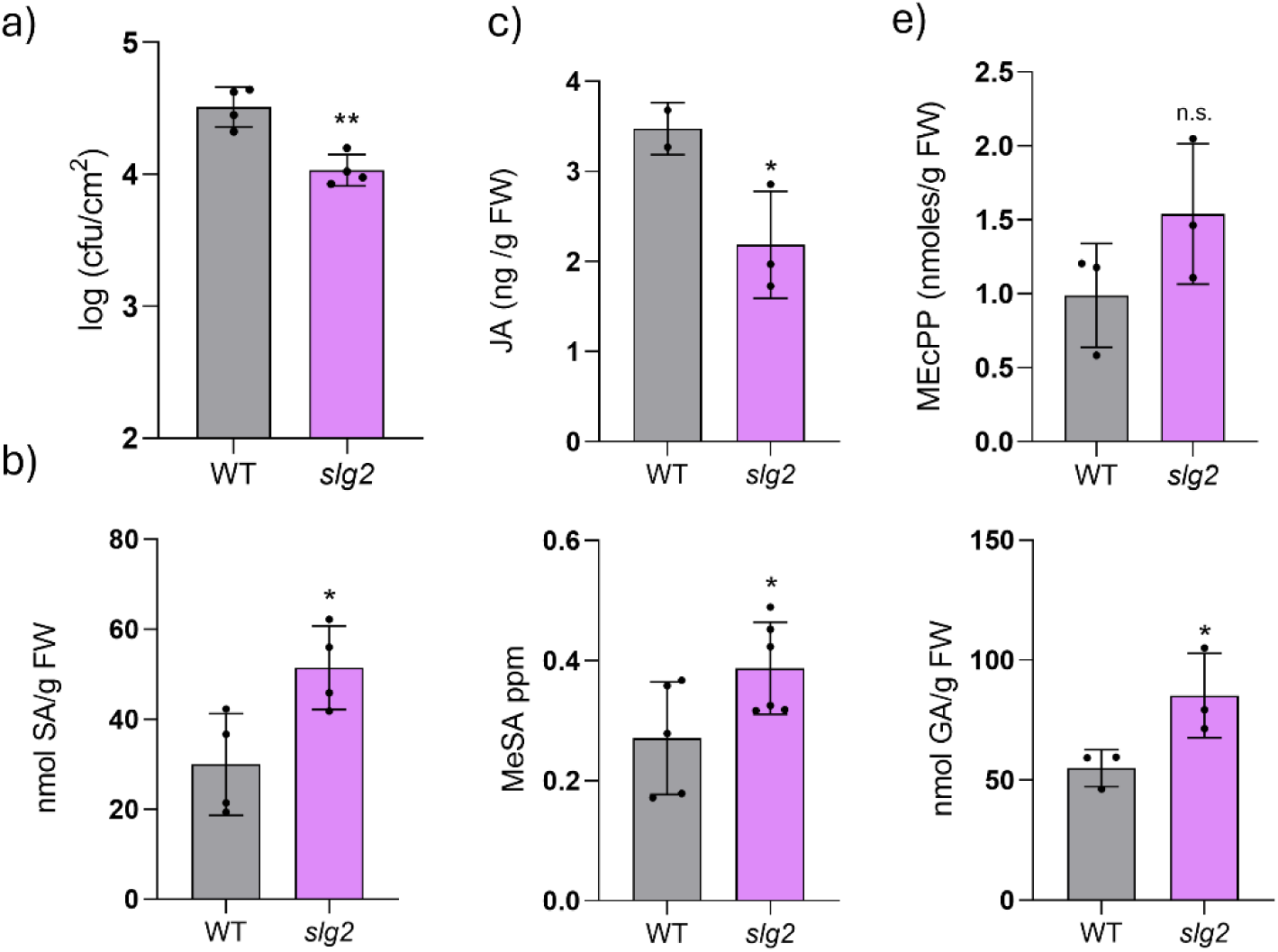
*Pst*-infected *sgl2* plants show higher SA levels and enhanced resistance to bacterial infection. (a) Bacterial content in *Pst*-infected WT and *slg2* plants. (b) Levels of total salicylic acid (SA) and derived metabolites methyl salycilate (MeSA) and gentisic acid (GA). (c) Levels of jasmonic acid (JA). (d) Levels of MEcPP. Bars and errors correspond to the mean and SD of n=>3 plants. Significant differences between WT and *slg2* values (t test) are shown with asterisks (* p < 0.05; ** p < 0.01); ’n.s.’ denotes no significant difference.

The activation of SA biosynthesis is essential for the defense response against *Pst* infection (Robert-Seilaniantz et al. 2011). Therefore, the levels of this defensive hormone and its derivatives gentisic acid (GA) and methyl salicylate (MeSA) were analyzed in WT and mutant *slg2* plants infected with *Pst*. Additionally, levels of jasmonic acid (JA) were also quantified, since a cross-talk between SA and JA has been classically described, being manifested as a reciprocal antagonism (Thaler *et al*., 2012). Consistent with the enhanced resistance observed, SA, GA, and MeSA levels were significantly higher in the *slg2* mutants compared to the WT (Fig. 5b), while JA levels were lower in the mutants (Fig. 5c), further supporting the notion of antagonistic regulation between these two pathways.

SA is a phytohormone that activates plant defense pathways (Klessig et al., 2018). It can be synthesized in plants from phenylalanine via the phenylpropanoid route or the isochorismate pathway, with the latter playing a predominant role in biotic stress responses, accounting for 90% of SA production under these conditions (Wildermuth et al., 2001). Previous studies demonstrated that enhanced production of monoterpenoids in tomato transgenic lines overexpressing a monoterpene synthase resulted in reduced levels of the MEP intermediate MEcPP, an inducer of the isochorismate synthase (*ICS*) gene required for SA biosynthesis (Pérez-Pérez et al., 2024) (Fig. 3). By contrast, in this study, MEcPP levels (Fig. 5d) and *ICS* expression (Fig. 3) were comparable between WT and *slg2* plants, indicating that the loss of SlG2 does not alter this regulatory mechanism.

Collectively, our results indicate that loss of SlG2 activity promotes the tomato plant immune response against *Pst* by promoting the biosynthesis of HMTPs and subsequently SA.

### VOCs associated to the ‘death aroma’ upon *Pst* infection are reduced in *slg2* mutants

Given that the loss of SlG2 activity results in an enhanced immune response, it is reasonable to expect that the increased resistance to bacterial infection observed in *slg2* plants would also be reflected in its VOC profile. To test this hypothesis, we first characterized the VOC profile of the compatible *Pst*-MicroTom interaction by comparing the volatilome of *Pst*-infected WT leaves with that of non-infected plants and then we compared the volatilome of *Pst*-infected WT and *slg2* mutant plants (Fig. 6).

**Fig. 6.**
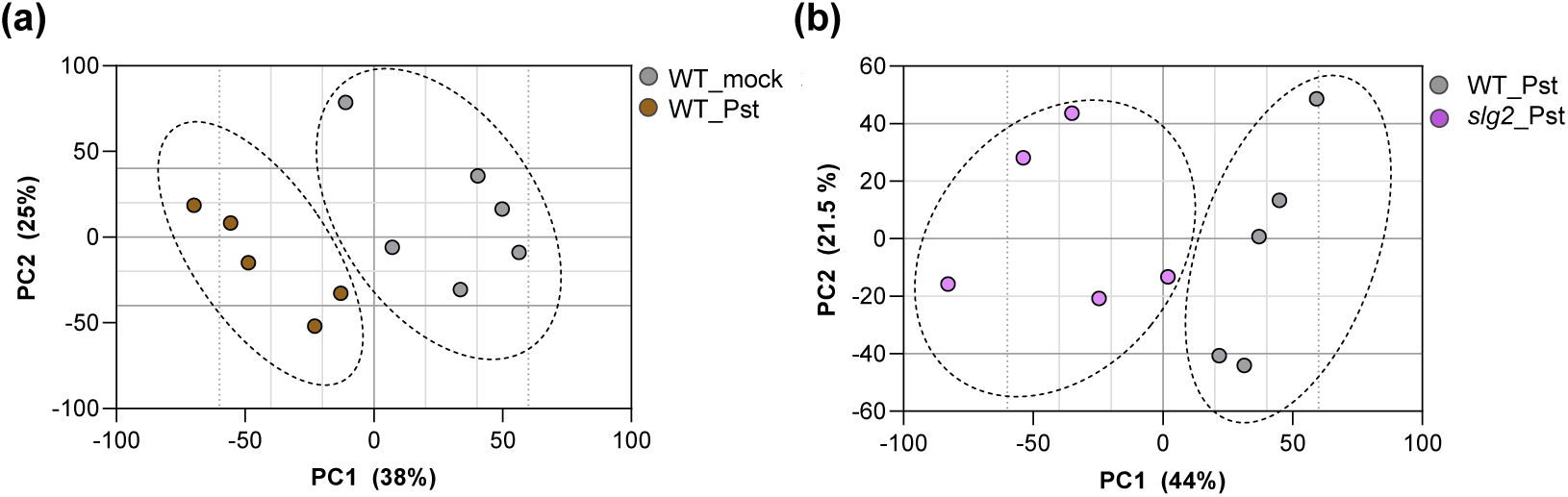
*Pst* infection alters the volatilome of MicroTom plants. Dots represent replicates for the indicated WT and *slg2* samples from leaves infected with *Pst* or treated with a mock solution without bacteria. Score plot of the PCA was based on the whole array of the mass spectra within a m/z range from 35 to 250. The dotted ellipses represent the 95% confidence intervals for each group.

In terms of the specific response of the MicroTom variety to *Pst* infection (Fig. 6a), the PCA revealed a clear separation between mock and *Pst*-infected plants along PC1 (38% of the variance), indicating substantial changes in VOC composition upon bacterial infection. The analysis of negative PC1 loading plots showed the differential compounds accumulated in MicroTom plants upon bacterial infection (Table 2). They were all unequivocally confirmed by using pure standards, except one of them, which was tentatively identified based on their mass spectra similarity (match > 900), and were named as components of the “scent of death”, which is based mainly on acid derivatives such as methyl benzoate (MeBA) as well as aldehyde and ketones derivatives including 1- penten-3-one, 3-methyl-2-butenal, 2-pentenal, 3-hexenal, and 4-oxohex-2-enal.

**Table 2.** List of differentially emitted VOCs in *Pst*-infected compared to mock-treated WT leaves. Only those showing a fold change (FC) >1.5 and p-values < 0.05 are shown. Family: KET, ketones; FUR, furanic; ALD, aldehydes; EST, esters. RT, retention time. Ion, major ion fragments used for quantification. Asterisk marks tentative identity.

| WT inf vs WT mock | VOC family | Ion | R.T (min) | Fold Change |
| --- | --- | --- | --- | --- |
| 1-Penten-3-one | KET | 55 | 11.26 | 1.92 |
| 2-Ethyl-Furan | FUR | 81 | 11.88 | 1.56 |
| 3-Methyl-2-Butenal | ALD | 84 | 13.58 | 2.29 |
| 2-Pentenal | ALD | 83 | 14.03 | 1.71 |
| 3-Hexenal | ALD | 80 | 15.74 | 1.95 |
| 4-Oxohex-2-enal * | ALD | 83 | 22.14 | 1.97 |
| Methyl Benzoate | EST | 77 | 27.20 | 28.65 |

Moreover, PCA comparing infected WT and *slg2* plants showed distinct clustering along PC1 (44% of the variance), reflecting the altered VOC profile expected in *slg2* mutants (Fig. 6b). In particular, the accumulation of VOCs forming the “death aroma” profile upon *Pst* infection of WT leaves (Table 2) was notably attenuated and even abolished in *slg2* mutants (Fig. 7). This reduced induction of cell death-associated VOCs in infected *slg2* mutants is consistent with their improved resistance to *Pst* infection. These results together demonstrate that both loss of SlG2 activity and *Pst* infection induce significant shifts in VOC profiles, with *slg2* mutants displaying a unique metabolic signature potentially linked to their enhanced resistance to bacterial infection.

**Fig. 7.**
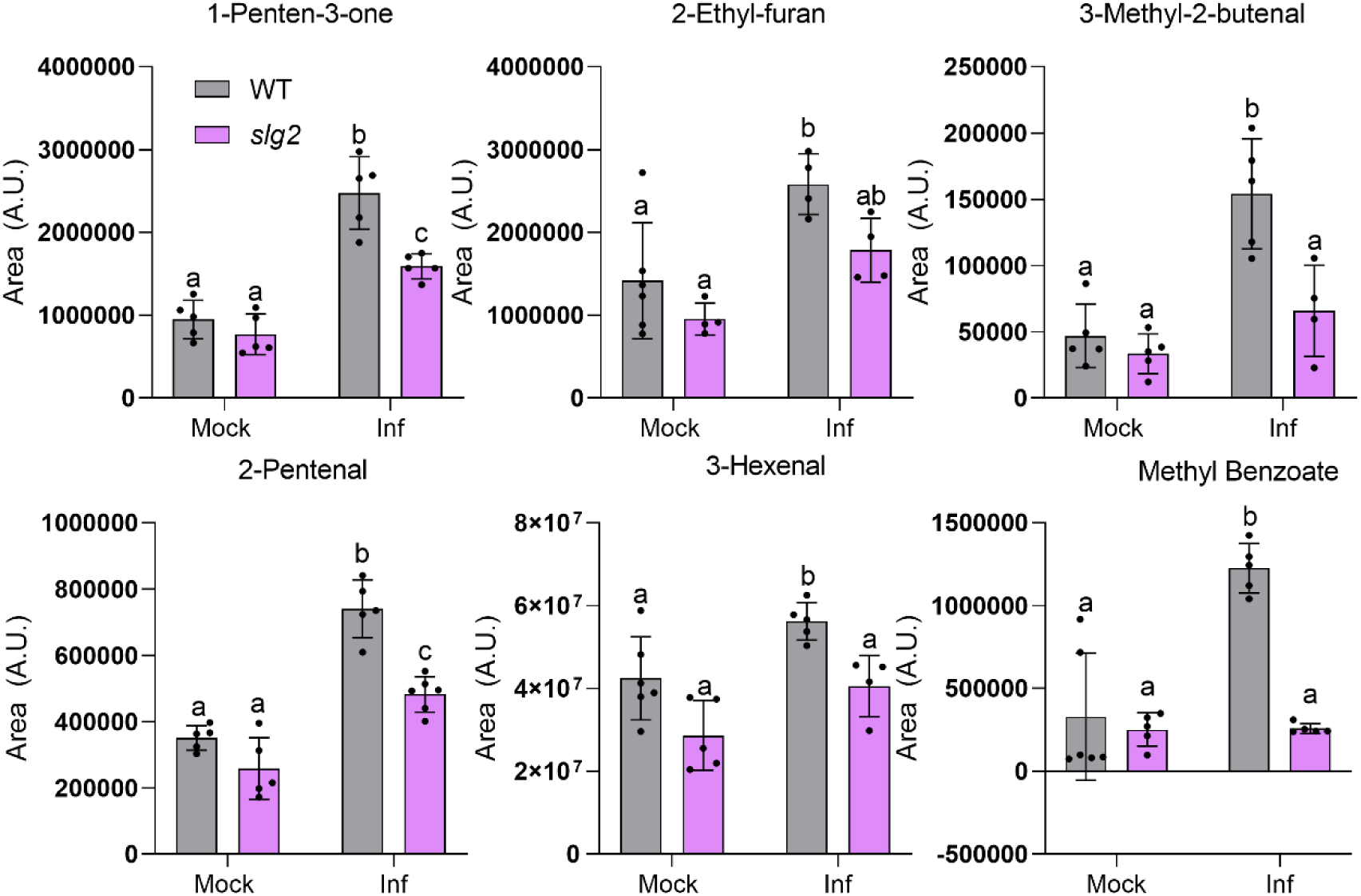
Mutant *sgl2* leaves release lower levels of death-associated VOCs. WT and mutant *slg2* leaves were infected with *Pst* or a mock solution without bacteria and samples were collected 24h later. Values represent the mean and SD of n=5 independent biological replicates. Different letters indicate statistically significant differences (two-way ANOVA with Dunnett’s multiple comparisons test).

## DISCUSSION

Our study provides novel insights into the role of the individual tomato GGPPS isoforms, with a major focus on the unexpected role of SlG2 isoform in regulating HMTP production and the associated enhancement of immune resistance responses in *slg2* plants. Our previous work using the same edited lines defective in individual GGPPS isoforms revealed that SlG1 and SlG3, but not SlG2, are involved in the production of GGPP-derived diterpenes such as geranyllinalool (GL) and its C16-homoterpene derivative (E,E)- 4,8,12-trimethyltrideca-1,3,7,11-tetraene (TMTT) (Ezquerro et al., 2023). This VOC is known to be released in response to biotic challenge (Ament et al., 2006; Falara et al., 2014). Under normal growth conditions, GL and TMTT levels were similar in WT and mutants of the three isoforms. However, following *Pst* infection, the induced increase was highly attenuated in *slg1* and *slg3* lines compared to WT and *slg2* plants (Ezquerro et al., 2023). Similar to TMTT, HMTPs have a defensive role, often acting as stress-responsive signals that provide protection to neighboring plants (Perez-Perez et al., 2024; Yu et al., 2024; Riedlmeier et al., 2017). Interestingly, in *slg2* plants, the levels of HMTPs were constitutively higher than those in WT, *slg1* and *slg3* plants prior to infection (Fig. 2), suggesting that they might also have a self-protective mechanism directly defending the plants that produce them from bacterial attack.

The most straightforward explanation for the mechanism by which the loss of SlG2 activity causes monoterpene overaccumulation is that removing SlG2 function results in a higher GPP supply for downstream monoterpene production. Expression of genes potentially involved in GPP synthesis such as *GPPS* and *SSU-I* remains similar in WT and *slg2* plants (Fig. 3), suggesting a mechanism independent of biosynthetic gene expression. One possibility might be that reduced consumption of IPP and DMAPP for GGPP biosynthesis when SlG2 is missing might divert the extra precursors towards GPP synthesis (Fig. 3). However, loss of other tomato GGPPS paralogs (particularly SlG3, which is the main GGPP-producing isoform) does not result in enhanced levels of GPP-derived monoterpenes (Fig. 2), suggesting that SlG2 might be specifically involved in GPP synthesis. *In vitro* studies have shown that heterodimerization with tomato SSU-I changes the product specificity of SlG1, SlG2 and SlG3 from GGPP to GPP, whereas heterodimerization with the SSU-II protein resulted in an improved specificity towards GGPP (Zhou and Pichersky 2020). Although the affinity of individual GGPPS isoforms for SSU-I or SSU-II proteins is still unknown, our *in silico* analyses suggested that SSU-I might preferentially bind to SlG2, resulting in more stable heterodimers compared to those potentially formed with SlG1 or SlG3 (Fig. 4). At the experimental level, it has been demonstrated that the SlG2/SSU-I heterodimer provides GPP for monoterpene biosynthesis in tomato fruit (Hivert et al., 2020). We speculate that SlG2/SSU-I heterodimers might also be the primary producers of GPP for monoterpene biosynthesis in leaves under normal conditions, whereas the other GGPPS isoforms (SlG1 and, predominatly, SlG3) might be primarily devoted to GGPP synthesis, at least in part by heterodimerizing with SSU-II subunits. This would explain why removal of SlG1 or SlG3 causes a reduction in GGPP-derived compounds such as diterpenes and carotenoids, respectively (Barja et al., 2021; Ezquerro et al., 2023; Burbano-Erazo et al., 2024) but no changes in GPP-derived monoterpenes (Fig. 2).

If SlG2/SSU-I heterodimers synthesize GPP in leaves, it might be expected that loss of SlG2 should result in lower GPP production. However, *slg2* mutant plants show increased (rather than decreased) accumulation of GPP-derived monoterpenes (Fig. 2). A possible explanation of this apparent contradiction was provided by the *in silico* analyses of the interaction of SSU-I with SlG1, SlG2 and SlG3 (Fig. 4 and Supplemental Fig. S1). It was concluded that SlG1/SSU-I and SlG3/SSU-I heterodimers might be more efficient at producing GPP than SlG2/SSU-I enzymes. In the absence of SlG2 proteins (i.e., in *slg2* plants), SSU-I monomers would become available to form heterodimers with SlG1 and/or SlG3, hence resulting in a higher production of GPP to be used for monoterpenes and HMTPs in *slg2* plants. These findings reinforce the hypothesis that plastidial prenyltransferase heterodimers have evolved distinct roles in the regulation of terpene biosynthesis, influenced by their protein-protein interactions and substrate preferences (Song et al., 2023).

Interestingly, the elevated levels of HMTPs such as α-terpineol in *slg2* plants resulted in a clear impact on defense mechanisms, confirming the defensive role of HMTPs in tomato (Pérez-Pérez et al., 2024). The *slg2* mutants exhibited significantly higher resistance to *Pst* infection compared to WT plants, as evidenced by a reduction in bacterial colony numbers (Fig. 5a). This improved resistance was correlated with increased levels of SA and its defense-related derivatives (MeSA and GA) in *slg2* leaves following *Pst* infection (Fig. 5b). SA is a critical phytohormone that is widely recognized as a signaling molecule that activates plant defense pathways (Klessig et al., 2018) but also its derivatives are related with an enhanced defensive response (Park et al.,2007; Bellés et al., 2006). Strikingly, increased HMTP levels in transgenic tomato plants constitutively overexpressing the monoterpene synthase gene *MTS1* (van Schie et al., 2007) did not result in increased *Pst* resistance or SA accumulation (Perez-Perez et al., 2024). It was proposed that enhanced consumption of GPP by MTS1 had a pull effect on the MEP pathway, resulting in decreased levels of intermediates such as MEcPP (Fig. 3). MEcPP is a well-characterized signal inducing the expression of the *ICS* gene involved in SA biosynthesis (Fig. 3) (Xiao et al., 2012; Pérez-Pérez et al., 2024). Decreased MEcPP levels in these *MTS1-*overexpressing and HMTP-accumulating transgenic lines hence reduced *ICS* expression and prevented SA synthesis (Gil et al., 2005; Pérez-Pérez et al., 2024). By contrast, *slg2* leaves show WT levels of MEcPP (Fig. 5d) and *ICS* expression (Fig. 3). This could be explained by our model on the mode of action of SlG2 presented here. If the increased GPP (and hence HMTP) production in the *slg2* mutant derives from the formation of novel SlG3/SSU-I heterodimers, it would be expected that the amount of remaining SlG3 proteins to produce GGPP via SlG3/SlG3 homodimers or/and SlG3/SSU- II heterodimers would decrease. In agreement with a reduced in GGPP synthesis in *slg2* mutants, they present reduced levels of GGPP-derived isoprenoids such as carotenoids, although the decrease is not as strong as in *slg3* plants (Barja et al., 2021; Burbano-Erazo et al., 2024). The lower GGPP production in the *slg2* mutant might compensate the increased production of GPP, resulting in no extra consumption of MEP pathway products (IPP and DMAPP) and hence no changes in the levels of intermediates, including MEcPP. The mechanism by which HMTPs regulate SA biosynthesis independently of the MEcPP / ICS pathway remains to be investigated. In any case, it could be another mechanism of metabolic regulation, where metabolite biosynthesis is connected with plant defense pathways, facilitating the balance between growth and defense under varying environmental conditions. Interestingly, the role of monoterpenes in the defensive response against *Pst* is gaining increasing attention, with a growing number of studies moving forward from the traditional context of plant-insect interaction responses to focus on the interaction with *Pst* (Pérez-Pérez et al., 2024; Riedlmeier et al., 2017; Wenig et al., 2019).

Our previous work with the Rio Grande cultivar showed that HMTPs were part of the VOC profile of tomato leaves infected with avirulent *Pst* strains, which defines an immunized plant response (López-Gresa et al. 2017; Perez-Perez et al., 2024). By contrast, the same tomato cultivar infected with virulent bacteria strains produced a VOC profile characterized by SA derivatives such as MeSA and salicylaldehyde, isoprenoid chlorides, sesquiterpenes, and monoterpenes such as α-pinene, α-phellandrene, β-phellandrene and limonene (López-Gresa et al. 2017). A different VOC profile was observed here in MicroTom plants. The absence of the *Pto* resistance gene in MicroTom implies a compatible interaction inducing the plant disease. Indeed, in response to *Pst* infection, MicroTom plants exhibited significant changes in the volatilome, with increased emission of methyl benzoate (MeBA), and several aldehyde and ketone derivatives, known components of the “death aroma” (Table 2 and Fig. 7). This response was also observed in *slg2* mutants, but the induction of these VOCs was notably reduced compared to the WT (Fig. 7). In rice, insect damage also triggers the production and emission of benzoic acid derivatives such as MeBA, which, along with MeSA contribute to defense signaling. The emission of MeBA in rice is tightly regulated by interactions between JA and SA signaling pathways, underscoring the multifaceted roles of benzoic acid derivatives in plant defense and inter-plant communication (Zhao et al., 2010). In turn, the role of specific aldehydes, such as 2-pentenal, has been highlighted in other plant-pathogen interactions. For instance, in resistant wheat cultivars inoculated with WSMV at elevated temperatures, 2-pentenal showed a significant increase in abundance, suggesting its potential role in mediating heat stress-specific defense responses (Farahbakhsh et al., 2023). This observation aligns with the idea that certain aldehydes can play a dual role, both as markers of tissue damage and as components of a coordinated defense mechanism. The attenuation of the infection-associated “scent of death” in the tomato *slg2* mutant plants is consistent with their enhanced resistance to *Pst* infection. The reduction of some death-associated VOCs likely reflects an overall decrease in damaged tissue, contributing to the ability of the plant to limit bacterial spread and maintain higher resistance.

Collectively, our results demonstrate that plastidial GGPPS isoforms are important contributors to the defense response of tomato plants against bacterial infection in terms of VOC production. While SlG1 and SlG3 are required to produce GGPP-derived defensive diterpenes, SlG2 regulates GPP-derived monoterpene (including HMTP) biosynthesis. Our work further provides strong experimental evidence connecting HMTPs and immune responses via SA in tomato plants. The loss of SlG2 enhances constitutive HMTP production, increases SA levels and attenuates the release of cell death-associated VOCs upon *Pst* infection, ultimately improving the plant resistance to bacteria. These findings provide valuable insights into the intricate regulation of VOC production and defense signaling pathways in plants and may have broader implications for developing crop varieties with enhanced resistance to pathogens.

## ACKNOWLEDGEMENTS

We would like to thank the IBMCP Metabolomics Platform (Valencia, Spain), especially Teresa Caballero for her support in VOC injections, Ana Espinosa for MEcPP quantification and Esther Carrera and Jorge Baños for JA measurements. This work was supported by grants from Spanish MCIN/AEI/10.13039/501100011033 and European NextGeneration EU/PRTR and PRIMA programs (PID2023-149584NB-I00, PID2023- 152361OB-I00, PID2020-115810GB-I00, PID2020-116765RB-I00 and UToPIQ-PCI2021-121941) as well as by Generalitat Valenciana (PROMETEU/2021/056 and AGROALNEXT/2022/067). M.R.-C. lab is part of the MCIN/AEI-funded Spanish Carotenoid Network, CaRed (RED2022-134577-T) and the EU/COST-funded ReCrop network (Reproductive Enhancement of CROP resilience to extreme climate, CA22157). S.-H.H. lab is supported by the National Research Foundation of Korean grant (RS-2024- 00347806) funded by the Ministry of Science and ICT, South Korea. J.P.-P. was a recipient of a JAEINT_21_02081 of the Consejo Superior de Investigaciones Científicas and currently holds a predoctoral contract of the Ministerio de Universidades (FPU21/00259).

M.E. received a predoctoral fellowship from MCIN/AEI (BES-2017-080652).

## AUTHOR CONTRIBUTIONS

M.P.L.-G. M.R.-C. and P.L. designed the research. J.P.-P., and M.E. performed the experimental research. S.L. and S.-H.H. performed the computational simulations. J.P.-P., S.-H.H., M.P.L.-G., M.R.-C. and P.L. analyzed and discussed the data. J.P.-P., M.P.L.-G., M.R.C. and P.L. wrote the paper.

## DATA AVAILABILITY

All data is incorporated into the article.

## SUPPLEMENTAL INFORMATION

**Supplementary Fig. S1.**
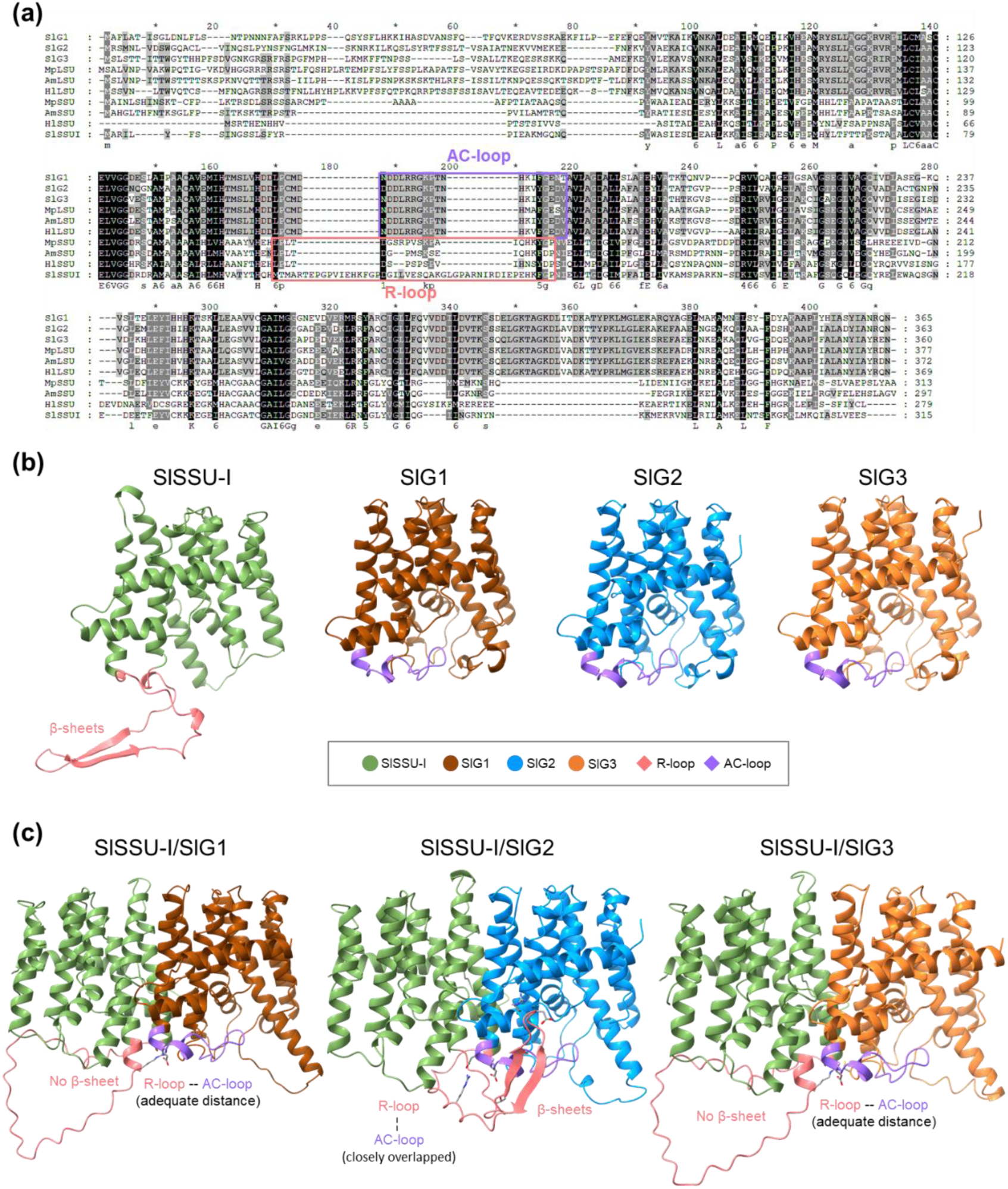
Sequence alignment and protein structure prediction. (a) Amino acid sequence alignment: SlG1 (*Solanum lycopersicum* GGPPS1), SlG2 (*S. lycopersicum* GGPPS2), SlG3 (*S. lycopersicum* GGPPS3), MpLSU (*Mentha piperita* LSU), AmLSU (*Antrirrhinum majus* LSU), HlLSU (*Humulus lupulus* LSU), MpSSU (*M. piperita* SSU), AmSSU (*A. majus* SSU), HlSSU (*H. lupulus* SSU), SlSSUI (*S. lycopersicum* SSU-I). All sequences presented here have the N- terminal signal peptides omitted. (b) Predicted monomeric protein structures using AlphaFold3. (c) Predicted heterodimeric protein structures using AlphaFold3. All protein structures were visualized with ChimeraX.

**Supplementary Table S1.** PDBePISA analysis of interface summary between tomato SSU-I and SlG1, SlG2, or SlG3 in heterodimers.

| Interfacing value |  | SISSU-I/SIG1 heterodimer |  |  |  | SISSU-I/SIG2 heterodimer |  |  |  | SISSU-I/SIG3 heterodimer |  |  |  |
| --- | --- | --- | --- | --- | --- | --- | --- | --- | --- | --- | --- | --- | --- |
|  |  | SISSU-I |  | SIG1 |  | SISSU-I |  | SIG2 |  | SISSU-I |  | SIG3 |  |
| Number of atoms | Interface | 179 | 7.9% | 178 | 8.1% | **202** | **9.0%** | **214** | **9.8%** | 179 | 7.9% | 181 | 8.1% |
|  | Surface | 1372 | 60.8% | 1253 | 56.8% | 1351 | 59.9% | 1252 | 57.3% | 1378 | 61.1% | 1292 | 57.6% |
|  | Total | 2255 | 100.0% | 2207 | 100.0% | 2255 | 100.0% | 2186 | 100.0% | 2255 | 100.0% | 2244 | 100.0% |
| Number of residues | Interface | 50 | 17.2% | 50 | 17.2% | **56** | **19.2%** | **62** | **21.1%** | 49 | 16.8% | 47 | 15.9% |
|  | Surface | 259 | 89.0% | 263 | 90.4% | 261 | 89.7% | 266 | 90.5% | 260 | 89.3% | 256 | 86.5% |
|  | Total | 291 | 100.0% | 291 | 100.0% | 291 | 100.0% | 294 | 100.0% | 291 | 100.0% | 296 | 100.0% |
| Solvent-accessible area (Å<sup>2</sup>) | Interface | 1939.7 | 11.2% | 1922.9 | 13.2% | **2165.6** | **13.1%** | **2195.5** | **15.3%** | 1972.2 | 11.5% | 1945.6 | 12.7% |
|  | Total | 17359.5 | 100.0% | 14557.1 | 100.0% | 16481.5 | 100.0% | 14396.2 | 100.0% | 17169.8 | 100.0% | 15323.8 | 100.0% |
| Solvation energy (kcal/mol) | Isolated structure | -244.6 | 100.0% | -278.6 | 100.0% | -249 | 100.0% | -275.8 | 100.0% | -246.2 | 100.0% | -281.4 | 100.0% |
|  | Gain on complex formation | -16 | 6.6% | -19.4 | 7.0% | **-21.4** | **8.6%** | **-21.6** | **7.8%** | -16.4 | 6.7% | -20.4 | 7.3% |
|  | Average gain | -6.6 | 2.7% | -4.8 | 1.7% | **-6.7** | **2.7%** | **-8.2** | **3.0%** | -6.4 | 2.6% | -5.8 | 2.1% |
|  | P-value | 0.021 |  | 0.000 |  | 0.001 |  | 0.002 |  | 0.015 |  | 0.001 |  |

**Suplementary Table S2.**
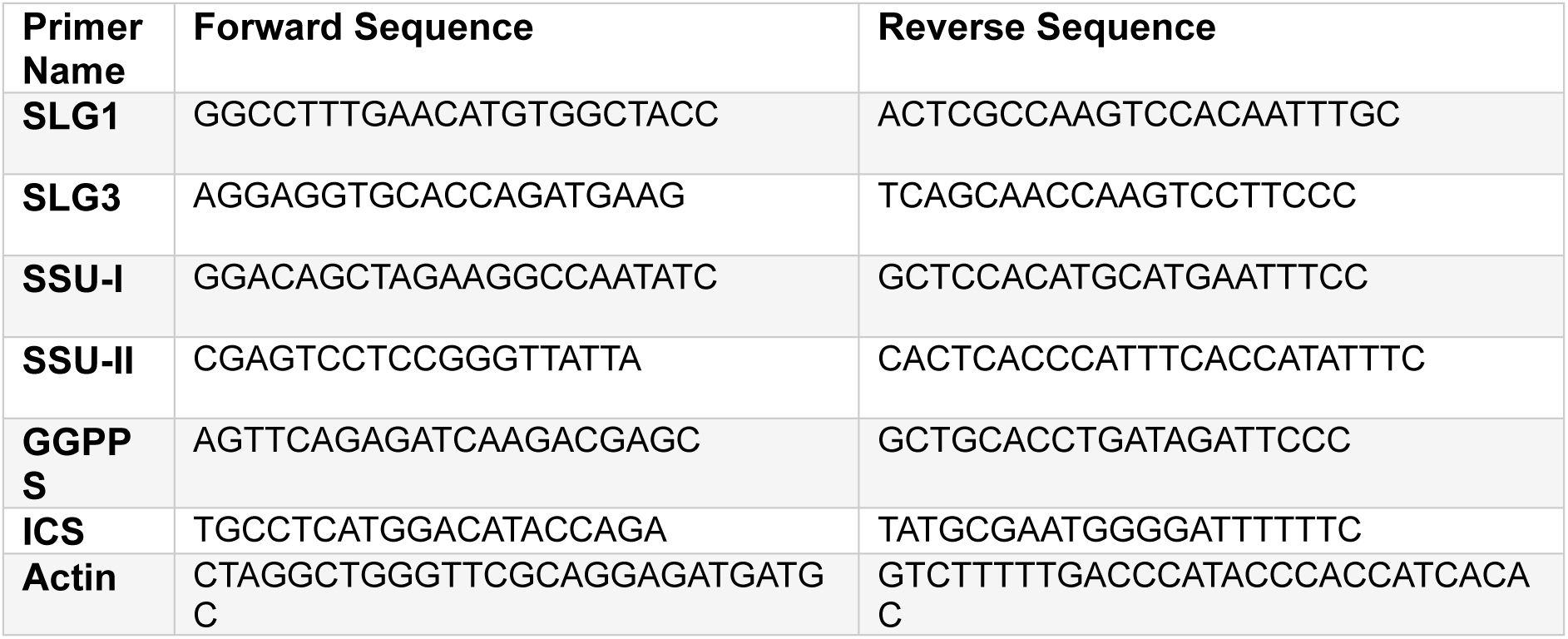
Listed qPCR primers used.

## Notes

### Competing Interest Statement

The authors have declared no competing interest.

## REFERENCES

Abramson J, et al. (2024). Accurate structure prediction of biomolecular interactions with AlphaFold 3. Nature 636: E4.

Ament K, Van Schie CC, Bouwmeester HJ, Haring MA, Schuurink RC (2006) Induction of a leaf specific geranylgeranyl pyrophosphate synthase and emission of (E,E)-4,8,12- trimethyltrideca-1,3,7,11-tetraene in tomato are dependent on both jasmonic acid and salicylic acid signaling pathways. Planta 224: 1197–1208

Baidoo EEK, Xiao Y, Dehesh K, Keasling JD.(2014) Metabolite profiling of plastidial deoxyxylulose- 5-phosphate pathway intermediates by liquid chromatography and mass spectrometry. Methods Mol Biol: 1153:57–76.

Barja MV, Ezquerro M, Beretta S, Diretto G, Florez-Sarasa I, Feixes E, Fiore A, Karlova R, Fernie AR, Beekwilder J, et al (2021) Several geranylgeranyl diphosphate synthase isoforms supply metabolic substrates for carotenoid biosynthesis in tomato. New Phytologist 231: 255–272

Barja MV, Rodriguez-Concepcion M (2021) Plant geranylgeranyl diphosphate synthases: every (gene) family has a story. aBIOTECH 2: 289–298

Bellés JM, Garro R, Pallás V, Fayos J, Rodrigo I, Conejero V (2006). Accumulation of gentisic acid as associated with systemic infections but not with the hypersensitive response in plant-pathogen interactions. Planta 223:500–551

Burbano-Erazo E, Ezquerro M, Sanchez-Bel P, Rodriguez-Concepcion M (2025). Specific sets of geranylgeranyl diphosphate synthases and phytoene synthases control the production of carotenoids and ABA in different tomato tissues. Physiol Plant 177: e70052.

Campos L, Granell P, Tarraga S, Lopez-Gresa MP, Conejero V, Belles JM, Rodrigo I, Lison P. (2014). Salicylic acid and gentisic acid induce RNA silencing-related genes and plant resistance to RNA pathogens. Plant Physiol Biochem. 77:35–43.

Chang TH, Hsieh FL, Ko TP, Teng KH, Liang PH, Wang AH (2010). Structure of a heterotetrameric geranyl pyrophosphate synthase from mint (Mentha piperita) reveals intersubunit regulation. Plant Cell 22: 454–467.

Chen Q, Fan D, Wang G (2015) Heteromeric Geranyl(geranyl) Diphosphate Synthase Is Involved in Monoterpene Biosynthesis in Arabidopsis Flowers. Mol Plant 8: 1434–1437

Ezquerro M, Li C, Pérez-Pérez J, Burbano-Erazo E, Barja MV, Wang Y, Dong L, Lisón P, López-Gresa MP, Bouwmeester HJ, et al (2023) Tomato geranylgeranyl diphosphate synthase isoform 1 is involved in the stress-triggered production of diterpenes in leaves and strigolactones in roots. New Phytologist 239: 2292–2306

Falara V, Alba JM, Kant MR, Schuurink RC, Pichersky E (2014) Geranyllinalool Synthases in Solanaceae and Other Angiosperms Constitute an Ancient Branch of Diterpene Synthases Involved in the Synthesis of Defensive Compounds. Plant Physiol 166: 428–441

Farahbakhsh F, Massah A, Hamzehzarghani H, Yassaie M, Amjadi Z, El-Zaeddi H, Carbonell-Barrachina AA (2023) Comparative profiling of volatile organic compounds associated to temperature sensitive resistance to wheat streak mosaic virus (WSMV) in resistant and susceptible wheat cultivars at normal and elevated temperatures. 281: 153903. J Plant Physiol.

Gil MJ, Coego A, Mauch-Mani B, Jordá L, Vera P (2005). The Arabidopsis csb3 mutant reveals a regulatory link between salicylic acid-mediated disease resistance and the methyl-erythritol 4- phosphate pathway. Plant J 44:155–166.

Hivert G, Davidovich-Rikanati R, Bar E, Sitrit Y, Schaffer A, Dudareva N, Lewinsohn E (2020) Prenyltransferases catalyzing geranyldiphosphate formation in tomato fruit. Plant Science 296: 110504

Hsieh FL, Chang TH, Ko TP, Wang AH (2010). Enhanced specificity of mint geranyl pyrophosphate synthase by modifying the R-loop interactions. J Mol Biol 404: 859–873

Huey R, Morris GM, Forli S (2012). Using AutoDock 4 and AutoDock vina with AutoDockTools: a tutorial. The Scripps Research Institute Molecular Graphics Laboratory 10550.92037: 1000.

Klessig DF, Choi HW, Dempsey DA (2018) Systemic acquired resistance and salicylic acid: Past, present, and future. Molecular Plant-Microbe Interactions 31: 871–888

Krissinel E, Henrick K (2007). Inference of macromolecular assemblies from crystalline state. J Mol Biol 372:774–797.

López-Gresa MP, Lisón P, Campos L, Rodrigo I, Rambla JL, Granell A, Conejero V, Bellés JM (2017) A Non-targeted Metabolomics Approach Unravels the VOCs Associated with the Tomato Immune Response against Pseudomonas syringae. Front Plant Sci.8:1188

Ntoukakis V, Mucyn TS, Gimenez-Ibanez S, Chapman HC, Gutierrez JR, Balmuth AL, Jones AME, Rathjen JP (2009) Host Inhibition of a Bacterial Virulence Effector Triggers Immunity to Infection. Science 324: 784–787

Park SW, Kaimoyo E, Kumar D, Mosher S, Klessig DF (2007). Methyl salicylate is a critical mobile signal for plant systemic acquired resistance. Science:318:113–116

Pérez-Pérez J, Minguillón S, Kabbas-Piñango E, Payá C, Campos L, Rodríguez-Concepción M, Espinosa-Ruiz A, Rodrigo I, Bellés JM, López-Gresa MP, et al (2024) Metabolic crosstalk between hydroxylated monoterpenes and salicylic acid in tomato defense response against bacteria. Plant Physiol 195: 2323–2338

Pettersen EF, Goddard TD, Huang CC, Meng EC, Couch GS, Croll TI, Morris JH, Ferrin TE (2021). UCSF ChimeraX: Structure visualization for researchers, educators, and developers. Protein Sci 30:70–82.

Pravda L, Sehnal D, Toušek D, Navrátilová V, Bazgier V, Berka K, Svobodová Vareková R, Koca J, Otyepka M (2018). MOLEonline: a web-based tool for analyzing channels, tunnels and pores (2018 update). Nucleic Acids Res 46: 368–373.

Rambla JL, López-Gresa MP, Bellés JM, Granell A (2015) Metabolomic Profiling of Plant Tissues. 221–235

Riedlmeier M, Ghirardo A, Wenig M, Knappe C, Koch K, Georgii E, Dey S, Parker JE, Schnitzler JP, Vlot AC (2017) Monoterpenes support systemic acquired resistance within and between plants. Plant Cell 29: 1440–1459

Robert-Seilaniantz A, Grant M, Jones JDG (2011) Hormone crosstalk in plant disease and defense: More than just JASMONATE-SALICYLATE antagonism. Annu Rev Phytopathol 49: 317–343

Rodríguez-Concepción M, Boronat A (2015) Breaking new ground in the regulation of the early steps of plant isoprenoid biosynthesis. Curr Opin Plant Biol 25: 17–22

Seo M, Jikumaru Y, Kamiya Y (2011). “Profiling of hormones and related metabolites in seed dormancy and germination studies,” in Seed Dormancy. Methods and Protocols ed. Kermode A. R. (Totowa, NJ: Humana Press) 773: 99–111.

Song S, Jin R, Chen Y, He S, Li K, Tang Q, Wang Q, Wang L, Kong M, Dudareva N, Smith BJ, Zhou F, Lu S (2023) The functional evolution of architecturally different plant geranyl diphosphate synthases from geranylgeranyl diphosphate synthase. Plant Cell 35: 2293–2315.

van Schie CCN, Haring MA, Schuurink RC (2007) Tomato linalool synthase is induced in trichomes by jasmonic acid. Plant Mol Biol 64: 251–263

Thaler JS, Humphrey PT, Whiteman NK (2012) Evolution of jasmonate and salicylate signal crosstalk. Trends Plant Sci 17: 260–270

Tholl D, Kish CM, Orlova I, Sherman D, Gershenzon J, Pichersky E, Dudareva N (2004) Formation of Monoterpenes in *Antirrhinum majus* and *Clarkia breweri* Flowers Involves Heterodimeric Geranyl Diphosphate Synthases. Plant Cell 16: 977–992

Vázquez Prol F, Márquez-Molins J, Rodrigo I, López-Gresa MP, Bellés JM, Gómez G, Pallás V, Lisón P (2021) Symptom Severity, Infection Progression and Plant Responses in Solanum Plants Caused by Three Pospiviroids Vary with the Inoculation Procedure. Int J Mol Sci 22: 6189

Vranová E, Coman D, Gruissem W (2013) Network Analysis of the MVA and MEP Pathways for Isoprenoid Synthesis. Annu Rev Plant Biol 64: 665–700

Wenig M, Ghirardo A, Sales JH, Pabst ES, Breitenbach HH, Antritter F, Weber B, Lange B, Lenk M, Cameron RK, et al (2019) Systemic acquired resistance networks amplify airborne defense cues. Nat Commun 10: 3813

Wildermuth MC, Dewdney J, Wu G, Ausubel FM (2001) Isochorismate synthase is required to synthesize salicylic acid for plant defence. Nature. 414: 562–565

Xiao Y, Savchenko T, Baidoo EEK, Chehab WE, Hayden DM, Tolstikov V, Corwin JA, Kliebenstein DJ, Keasling JD, Dehesh K (2012). Retrograde signaling by the plastidial metabolite MEcPP regulates expression of nuclear stress-response genes. Cell 149:1525–1535. 10.1016/j.cell.2012.04.038

Yu H, Buchholz A, Pullinen I, Saarela S, Li Z, Virtanen A, Blande JD (2024) Biogenic secondary organic aerosol participates in plant interactions and herbivory defense. Science. 385: 1225– 1230

Zhao N, Guan J, Ferrer JL, Engle N, Chern M, Ronald P, Tschaplinski TJ, Chen F (2010) Biosynthesis and emission of insect-induced methyl salicylate and methyl benzoate from rice. Plant Physiology and Biochemistry 48: 279–287

Zhou F, Pichersky E (2020) The complete functional characterisation of the terpene synthase family in tomato. New Phytologist 226: 1341–1360

